# Decoding Brain Dynamics Via Cross-Network Non-Linear Trajectories

**DOI:** 10.1101/2025.11.25.690593

**Authors:** Masoud Seraji, Sarah Shultz, Qiang Li, Liang Ma, Zening Fu, Vince D Calhoun

## Abstract

Infant brain networks mature rapidly and nonlinearly, yet cross-network comparisons of developmental trajectories remain rare. We introduce *cross-network non-linear trajectory characterization* (CNTC), which fits cubic polynomials to age-related change in three spatial measures—network-averaged spatial similarity (NASS), network strength, and network size—derived from resting-state fMRI. Using 137 scans from 74 neurotypical infants spanning 0–6 months of age, we estimated networks via group ICA, modeled age trajectories within each network, and then contrasted coefficients across networks. Across seven canonical systems, trajectories showed robust age effects. Cross-network differences were dominated by linear terms, with selective quadratic effects and few cubic effects. Visual and cerebellar systems exhibited the steepest NASS slopes and pronounced increases in strength and size, whereas motor/attention changes were more gradual. CNTC provides a compact, coefficient-level summary of inter-network maturation and a basis for benchmarking atypical development in future work.

## 1. INTRODUCTION

Infancy is a crucial period for brain maturation, during which fundamental functional networks emerge and reorganize rapidly [1]. Resting-state fMRI (rsfMRI) provides a noninvasive way to map these large-scale networks in sleeping infants [2]. Prior work has shown that canonical sensorimotor, visual, and association networks are already present at birth and that the pace of maturation differs across systems [3]. Because many developmental processes are non-linear, charting infant growth often requires flexible models [4]. Charting such trajectories may reveal early biomarkers of atypical development [2]. To date, however, most infant imaging studies have examined changes within individual networks or used simple spatial summaries, leaving cross-network developmental comparisons largely unexplored [5, 6].

In this study, we introduce cross-network nonlinear trajectory characterization (CNTC) to address this gap. CNTC fits a third-order (cubic) polynomial to each network-specific developmental metric as a function of age. The coefficients capture the linear and non-linear changes of the dynamics. Such polynomial modeling is well motivated by prior work showing that many cortical regions follow complex trajectories that often require quadratic or cubic fits [7]. By estimating coefficients for each metric and network, CNTC quantifies and compares the shape of developmental growth across systems [8].

We apply CNTC to three spatial network metrics (such as network-averaged spatial similarity (NASS), network strength, and network size) derived from functional networks in infant rsfMRI data. This network-to-network analysis goes beyond traditional single-network approaches. The novelty of our work is underscored by its contrast with prior analyses. Seraji et al. (2024) examined infant network development by measuring spatial features to characterize age-related reorganization [2]. That spatial analysis demonstrated how network maps consolidate with age, but it did not explicitly compare them across networks. In contrast, our CNTC framework explicitly fits linear, quadratic and cubic components to each network’s trajectory, enabling direct cross-system comparisons of developmental shape. Thus, to our knowledge, this is the first work to leverage polynomial trajectory modeling for inter-network developmental analysis in infancy, offering a novel perspective on how functional systems evolve relative to one another.

## 2. METHODS

### 2.1 Imaging Dataset

We analyzed longitudinal rs-fMRI from 74 neurotypically developing infants (43 males, 31 females) scanned between birth and 6 months. Infants had no familial autism, developmental delay, perinatal complications, neurological/sensory or genetic disorders, seizures, or MRI contraindications. Age was expressed as corrected age (adjusted to a 40-week gestational baseline) rather than chronological age to better index developmental stage [9]. Written parental consent was obtained; procedures were approved by Emory University IRB. Additional acquisition details appear in [2].

### 2.2 Preprocessing

The first 16 volumes were discarded to ensure magnetization steady state. Motion correction was performed with FSL `MCFLIRT`. Single-band reference images with opposite phase-encoding were used to estimate susceptibility-induced off-resonance fields and correct distortions in the multiband rs-fMRI, followed by slice-timing correction. Spatial normalization used a two-step procedure: (i) co-registration to the 3-month Baby Connectome Project T1 template [10] to respect developmental anatomy; (ii) alignment to MNI space using an EPI template [11]. Images were smoothed with a 6 mm FWHM Gaussian kernel.

### 2.3 Network Estimation

Quality control excluded scans with poor alignment to MNI space by comparing individual brain masks to a study-specific group mask (See [2]). Large-scale networks were identified with GIFT toolbox. For each subject, principal component analysis reduced dimensionality to 30 components, which were concatenated and further reduced at the group level to 20 PCs [12]. Group ICA (Infomax) was repeated 100 times to assess stability; the most reliable components (cluster-centroid alignment) were retained. Subject-specific maps were estimated using group-information guided ICA (GIG-ICA) [13]. Components matching canonical spatial motifs were labeled as 13 networks: primary/secondary visual, sub-cortical, cerebellum, primary/secondary motor, attention, default mode, temporal, auditory, and 3 frontal networks. We used 7 of them in this paper.

### 2.4 Spatial metrics

From each ICA spatial map, we computed three measures:

1. **Network-averaged spatial similarity (NASS):** Similarity between an individual map and the group template, computed as the Pearson correlation within the brain mask:

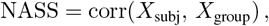

where larger values indicate closer alignment to the group-level spatial pattern.
2. **Network strength:** Mean contribution of suprathreshold voxels, indexing average engagement:

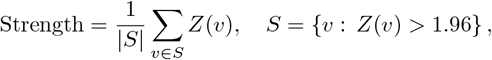

with *Z*>1.96 (two-sided *p*=0.05).
3. **Network size:** Spatial extent of the network, defined as the count of suprathreshold voxels:

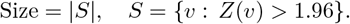

### 2.5 Cross-Network Nonlinear Trajectory Characterization

To characterize age-related change and compare it across networks, we fit a cubic polynomial to each metric within each network as a function of corrected age *t* (days):

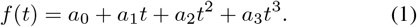

Coefficients *a*_1_, *a*_2_, and *a*_3_ respectively index linear (steady) change, quadratic curvature (acceleration/deceleration), and cubic inflection (S-shaped dynamics). Models were estimated by ordinary least squares, yielding coefficient estimates and their covariance.

#### Cross-network contrasts

For each polynomial term *k* ∈ {1, 2, 3} and every network pair (*i, j*), we formed 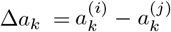 and computed a *z*-statistic using the pooled standard error from the coefficient covariance:

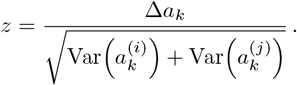

Two-sided *p*-values were obtained from the standard normal distribution (MATLAB), and multiple comparisons were controlled using Benjamini–Hochberg FDR (*q*<.05) across all network pairs, metrics, and polynomial terms. CNTC isolates where developmental trajectories differ in shape and magnitude across systems.

## 3. RESULTS

This section has two parts. First, we fit a cubic model to each metric within each of the seven networks to characterize within–network developmental trajectories (Fig. 1). Second, we compared polynomial coefficients across networks to identify where trajectories differ in shape and rate (Fig. 2).

**Fig. 1.**
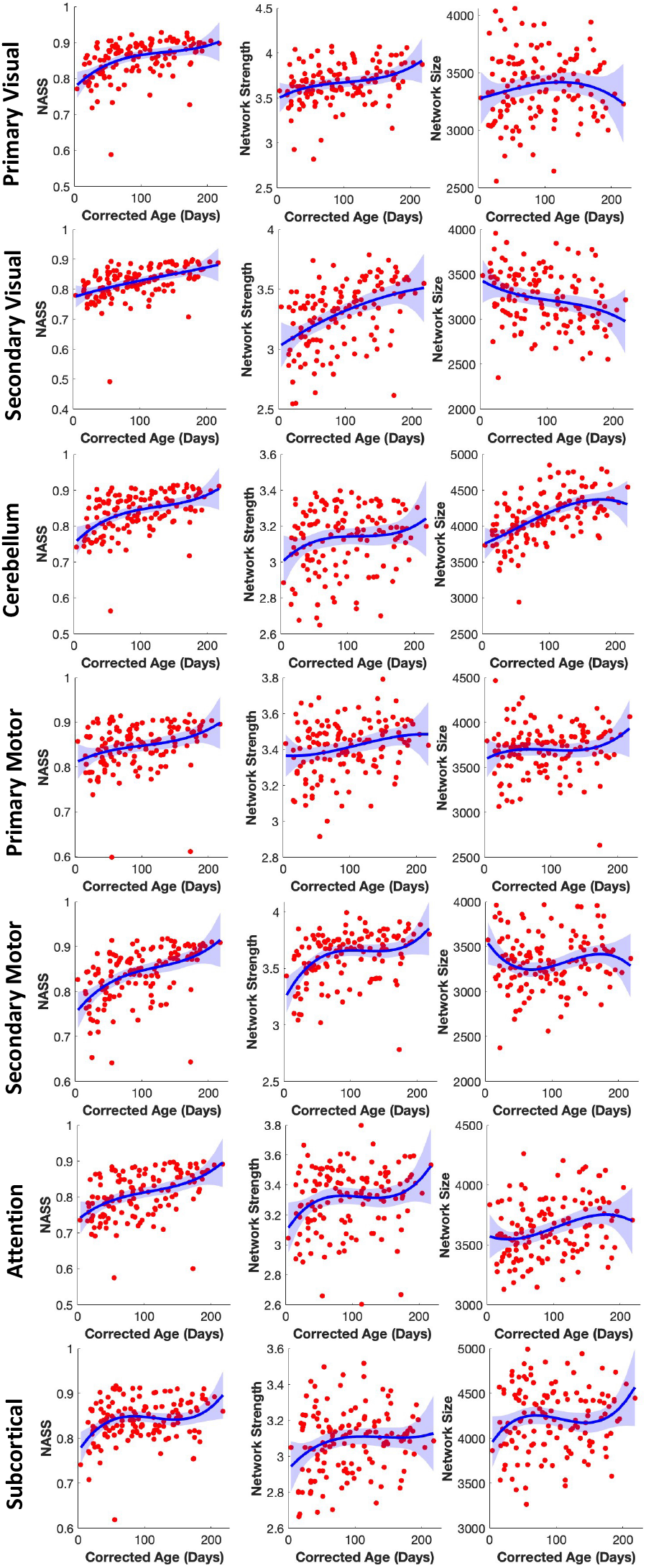
Non-linear Trajectory of Spatial Measurements. Cubic developmental trajectories of three spatial metrics across six infant brain networks.

**Fig. 2.**
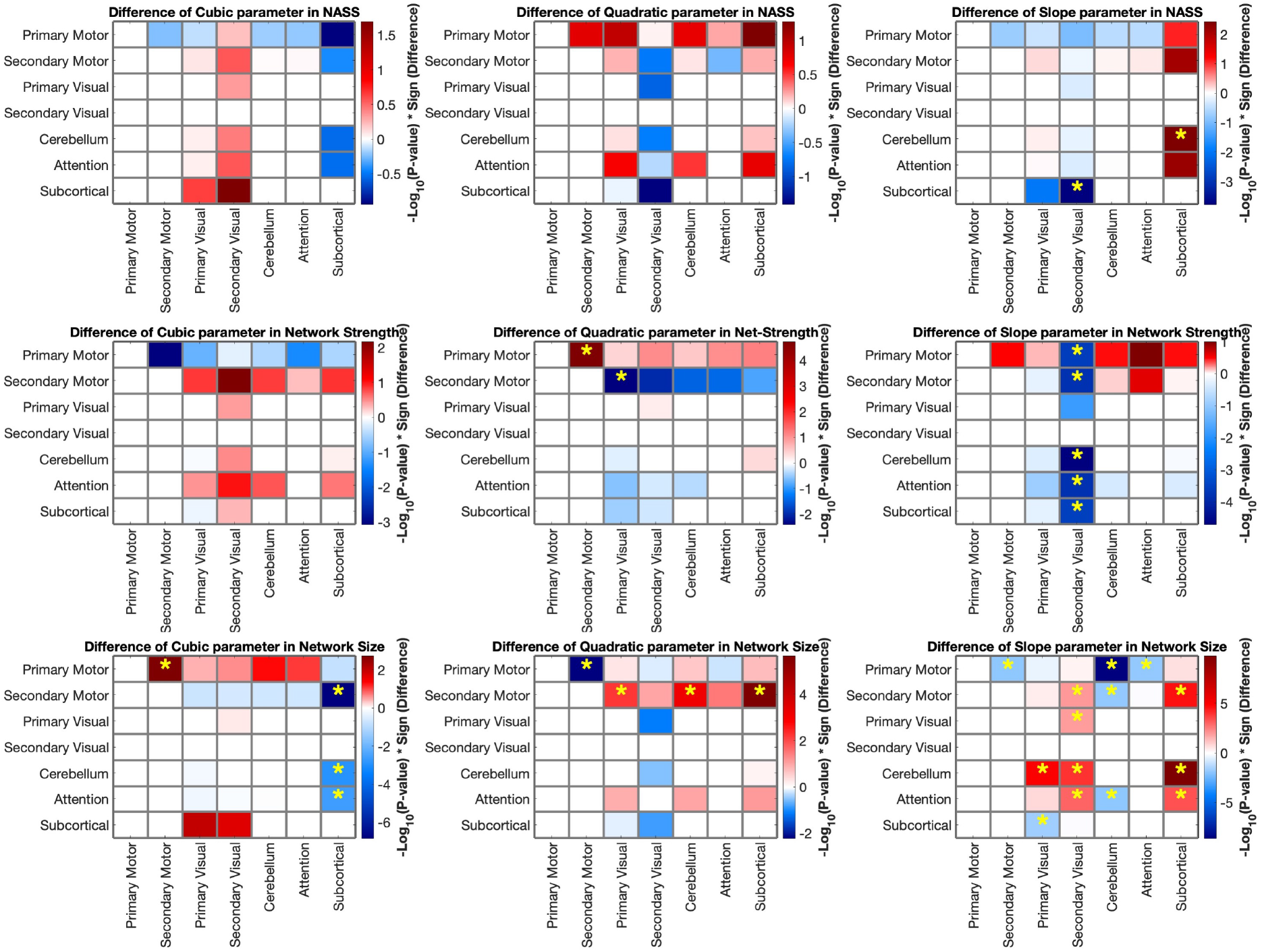
Cross-network coefficient contrasts. Heatmaps show pairwise differences in cubic, quadratic, and slope terms for three spatial metrics (NASS, network strength, network size). Values are − log_10_(*p*) × sign(Δ); yellow stars denote FDR-significant contrasts (*q* < 0.05). Rows/columns correspond to networks; blank cells on the diagonal are self-comparisons.

### 3.1 Nonlinear trajectories within networks

For each metric, trajectories were modeled as cubic polynomial. Fig. 1 shows the fitted curves for the primary/secondary visual, primary/secondary motor, cerebellum, attention and subcortical networks. Across metrics, networks exhibited age-related change with varying degrees of curvature. In general, quadratic components (*a*_2_) captured the dominant trend, with linear (*a*_1_) and cubic (*a*_3_) terms providing additional flexibility to accommodate accelerations and occasional inflection patterns. Visual inspection suggests earlier increases for visual systems and more gradual changes for motor/attention, but the extent and timing differ by metric. NASS increases in all six networks, with the steepest rise in the visual and secondary motor networks and a more gradual gain in primary motor and attention. Network strength exhibits modest growth overall, most evident in the visual and secondary motor networks, while the cerebellum shows a shallow increase. Network size expands with age for the cerebellum and primary visual network, whereas motor and attention are comparatively flat or weakly curved. Several trajectories display mid-infancy inflection points, consistent with nonlinear development, and uncertainty widens at the youngest/oldest ages due to fewer scans.

### 3.2 Cross-network coefficient contrasts

To quantify differences in trajectory shape, we contrasted coefficients between every network pair for each term *a*_1_, *a*_2_, *a*_3_ and metric, converting contrasts to two-sided *p*-values and controlling FDR at *q*<0.05. Fig. 2 summarizes coefficient-level contrasts between networks for the three spatial metrics. **NASS**. Cubic and quadratic terms showed no reliable cross-network differences. Significant effects were confined to the *slope* term, indicating that networks primarily differ in the *rate* at which individual maps converge to the group template rather than in higher-order trajectory shape.

#### Network Strength

Cubic differences were negligible. Several *quadratic* contrasts reached significance, and additional effects appeared in the *slope* term, suggesting metric-specific curvature plus rate differences in aggregate voxel engagement.

#### Network Size

This metric displayed the richest cross-network divergence. While cubic effects were limited to a few pairs, numerous *quadratic* and *slope* contrasts survived FDR, indicating heterogeneous growth in spatial extent across systems.

Overall, cross-network variability is dominated by differences in linear rate, with quadratic curvature contributing selectively (especially for size) and cubic inflections being rare.

## 4. DISCUSSION

Our analysis of infant resting-state fMRI revealed pronounced non-linear growth across networks and spatial metrics. In all seven networks, the fitted polynomial trajectories showed significant linear, quadratic, and often cubic components, reflecting complex maturation. Notably, the primary visual, secondary visual, and cerebellar networks exhibited the largest positive slopes and strong higher-order terms (Fig. 1), indicating rapid early maturation. In contrast, attention displayed slower, more protracted increases, with more pronounced nonlinear (quadratic/cubic) shapes. These slope differences in NASS imply distinct maturation rates: sensory/motor regions align with adult-like topology earlier, whereas integrative networks follow delayed, nonlinear trajectories. Similarly, network strength rose steeply in visual and motor systems, suggesting aggressive voxel contribution in these networks. Finally, network size changes were heterogenous: the primary visual and cerebellar systems expanded markedly, while some other networks changed little. In particular, subcortical regions showed early volumetric growth consistent with developing connectivity. These spatial expansion differences point to non-uniform functional development across the brain.

These findings align with known developmental hierarchies: Gao et al. (2015) reported that primary sensory networks are essentially “adult-like” at birth while association networks strengthen over the first years [14]. Likewise, Cao et al. (2017) emphasize a primary-to-higher-order progression in infant connectomics [6]. In line with these principles, our NASS slopes were steepest in primary visual/motor systems and shallower in higher-order networks. The rapid growth of the cerebellum is supported by recent work showing early cerebellocortical connectivity and strengthening of cerebellar networks through infancy [15]. Indeed, we observed robust cerebellar size and strength increases, consistent with its known rapid postnatal expansion [15, 2]. Moreover, Grotheer et al. (2022) found that myelination accelerates fastest in posterior/superior pathways during the first months [16], which may underlie our observation of nonlinear surges in visual and motor network measures. Subcortical development also fits these patterns: e.g. Alex et al. (2024) report the amygdala (a key subcortical node) matures first, with protracted thalamic growth [17], echoing our finding of early reorganization in subcortical networks. In summary, the strong gains in visual and cerebellar networks we observed are consistent with other studies highlighting rapid sensorimotor and cerebellar maturation in infancy [14, 15], while the more gradual, nonlinear trajectories of secondary and higher-order networks mirror prior reports of hierarchical connectome development [6, 16].

